# An Ancestry Based Approach for Detecting Interactions

**DOI:** 10.1101/036640

**Authors:** Danny S. Park, Itamar Eskin, Eun Yong Kang, Eric R. Gamazon, Celeste Eng, Christopher R. Gignoux, Joshua M. Galanter, Esteban Burchard, Chun J. Ye, Hugues Aschard, Eleazar Eskin, Eran Halperin, Noah Zaitlen

## Abstract

I

**Background:** Epistasis and gene-environment interactions are known to contribute significantly to variation of complex phenotypes in model organisms. However, their identification in human association studies remains challenging for myriad reasons. In the case of epistatic interactions, the large number of potential interacting sets of genes presents computational, multiple hypothesis correction, and other statistical power issues. In the case of gene-environment interactions, the lack of consistently measured environmental covariates in most disease studies precludes searching for interactions and creates difficulties for replicating studies.

**Results:** In this work, we develop a new statistical approach to address these issues that leverages genetic ancestry in admixed populations. We applied our method to gene expression and methylation data from African American and Latino admixed individuals respectively, identifying nine interactions that were significant at *p* < 5×10^−8^, we show that two of the interactions in methylation data replicate, and the remaining six are significantly enriched for low p-values (*p* < 1.8×10^−6^).

**Conclusion:** We show that genetic ancestry can be a useful proxy for unknown and unmeasured covariates in the search for interaction effects. These results have important implications for our understanding of the genetic architecture of complex traits.

## II Background

Genetic association studies in humans have focused primarily on the identification of additive SNP effects through marginal tests of association. There is growing evidence that both epistatic and gene-environment (*G*×*E*) interactions contribute significantly to phenotypic variation in humans and model organisms[1–5]. In addition to explaining additional components of missing heritability, interactions lend insights into biological pathways that regulate phenotypes and improve our understanding of their genetic architectures. However, identification of interactions in human studies has been complicated by the computational and multiple testing burden in the case of epistatic interactions, and the lack of consistently measured environmental covariates in the case of *G*×*E* interactions[6,7].

To overcome these challenges, we leverage the unique nature of genomes from recently admixed populations such as African Americans, Latinos, and Pacific Islanders. Admixed genomes are mosaics of different ancestral segments[8] and for each admixed individual it is possible to accurately estimate *θ*, the proportion of ancestry derived from each ancestral population (e.g. the fraction of European/African ancestry in African Americans)[9]. Ancestry has been previously leveraged to demonstrate that an array of environmental and biomedical covariates are correlated with *θ* [10–20] and we therefore consider its use as a surrogate for unmeasured and unknown environmental exposures. *θ* is also correlated with the genotypes of SNPs that are differentiated between the ancestral populations, suggesting that *θ* may be effectively used as a proxy for detecting multi-way epistatic interactions. Therefore, we propose a new SNP by *θ* test of interaction in order to detect evidence of interaction in admixed populations.

We first investigate the properties of our method through simulated genotypes and phenotypes of admixed populations. In our simulations we demonstrate that differential linkage-disequilibrium (LD) between ancestral populations can produce false positive SNP by *θ* interactions when local ancestry is ignored. To accommodate differential LD, we include local ancestry in our statistical model and demonstrate that this properly controls this confounding factor. We also show that our approach, the Ancestry Test of Interaction with Local Ancestry (AITL), is well-powered to detect *G*×*E* interactions when *θ* is correlated with the environmental covariates of interest and multi-way epistatic interactions. The power for detecting pairwise *G*×*G* interactions at highly differentiated SNPs is lower than direct interaction tests even after accounting for the additional multiple testing burden. However, the results of our simulations show that AITL is well powered to detect multi-way epistasis involving tens or hundreds of SNPs of small effects, not detectable by pairwise tests.

We first examined molecular phenotypes by applying our method to gene expression data from African Americans, as well as DNA methylation data from Latinos. Gene expression traits have previously been shown to have large-scale differences as a function of genetic ancestry[13]. Other molecular phenotypes, such as LDL levels, have also been shown to be associated with genetic ancestry [13,16,21–24]. For gene expression in particular, Price *et al*. showed that the effects of ancestry on expression are widespread and not restricted to a handful of genes. Additionally, molecular phenotypes are often used in deep phenotyping and Mendelian randomization studies and are thus directly relevant to elucidating disease biology[25,26].

We identified one genome-wide significant interaction (*p* < 5×10^−8^) associated with gene expression in the African Americans and eight significant interactions (*p* < 5×10^−8^) associated with methylation in the Latinos. Two of the eight interactions associated with DNA methylation in the Latinos also replicated and the remaining six were enriched for low p-values (*p* < 1.8×10^−6^). To demonstrate that our approach works in larger data sets we also applied AITL to asthma case-control data from Latinos and observed well-calibrated test statistics. Together, these results provide evidence for the existence of interactions regulating expression and methylation and show that our approach is statistically sound.

## III Results

### Simulated Data

To determine the utility of using *θ* as a proxy for unmeasured and unknown environmental covariates, we applied the AITL to simulated 2-way admixed individuals. We tested *θ*_1_, the proportion of ancestry from ancestral population 1, for interaction with simulated SNPs (see Simulation Framework). Power was computed over 1,000 simulations, assuming 10,000 SNPS being tested, and using a Bonferroni correction p-value cutoff of 5×10^−6^. We calculated the power using assumed interaction effect sizes (either *β*_*G*×*G*_ or *β*_*G*×*G*_) of 0.1, 0.2, 0.3, and 0.4 (see Simulation Framework). Although the few interactions reported for human traits and diseases have smaller effects in terms of the phenotypic variance they explain, we simulated large effects because genetic and environmental effect sizes in omics data, such as the expression and methylation data considered here, are known to be of larger magnitude. For example, some cis-eQTL SNPs explain up to 50% of the variance of gene expression [27]. However for most phenotypes, known interactions will explain a very small proportion of the phenotypic variance, mainly due to the fact that so few interactions have been identified and replicated[28].

### Power When Using *θ* as a Proxy for Highly Differentiated SNPs

To determine whether using *θ* as a proxy for highly differentiated SNPs is more powerful than testing all pairs of potentially interacting SNPs directly, we simulated two interacting SNPS in 1000 admixed individuals (see Simulation Framework). We then tested for an interaction using AITL by replacing the genotypes at the highly differentiated SNP with 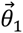. We observed that even with moderate effect sizes, using *θ* in place of the actual genotypes does not provide any increase in power even after accounting for multiple corrections (see Figure 1a). This is in agreement with recent work showing the limited utility of local ancestry by local ancestry interaction test to identify underlying SNP by SNP interaction when genotype data are available[29]. For the larger effect sizes we simulated, we do see power increasing as the delta between ancestral frequencies increases. The plots show that AITL has little power unless the effect was very strong. Figure 1b reveals that even with the multiple correction penalty, testing all pairwise SNPS directly is always more powerful. We note that when testing the interacting SNPs directly, we used a cutoff p-value of 1×10^−9^ since in theory we were testing all unique pairs of 10,000 SNPs. Based on these results, we would recommend testing for pairs of interacting SNPs directly if pairwise *G*×*G* interactions are a subject of interest in the study.

**Figure 1.**
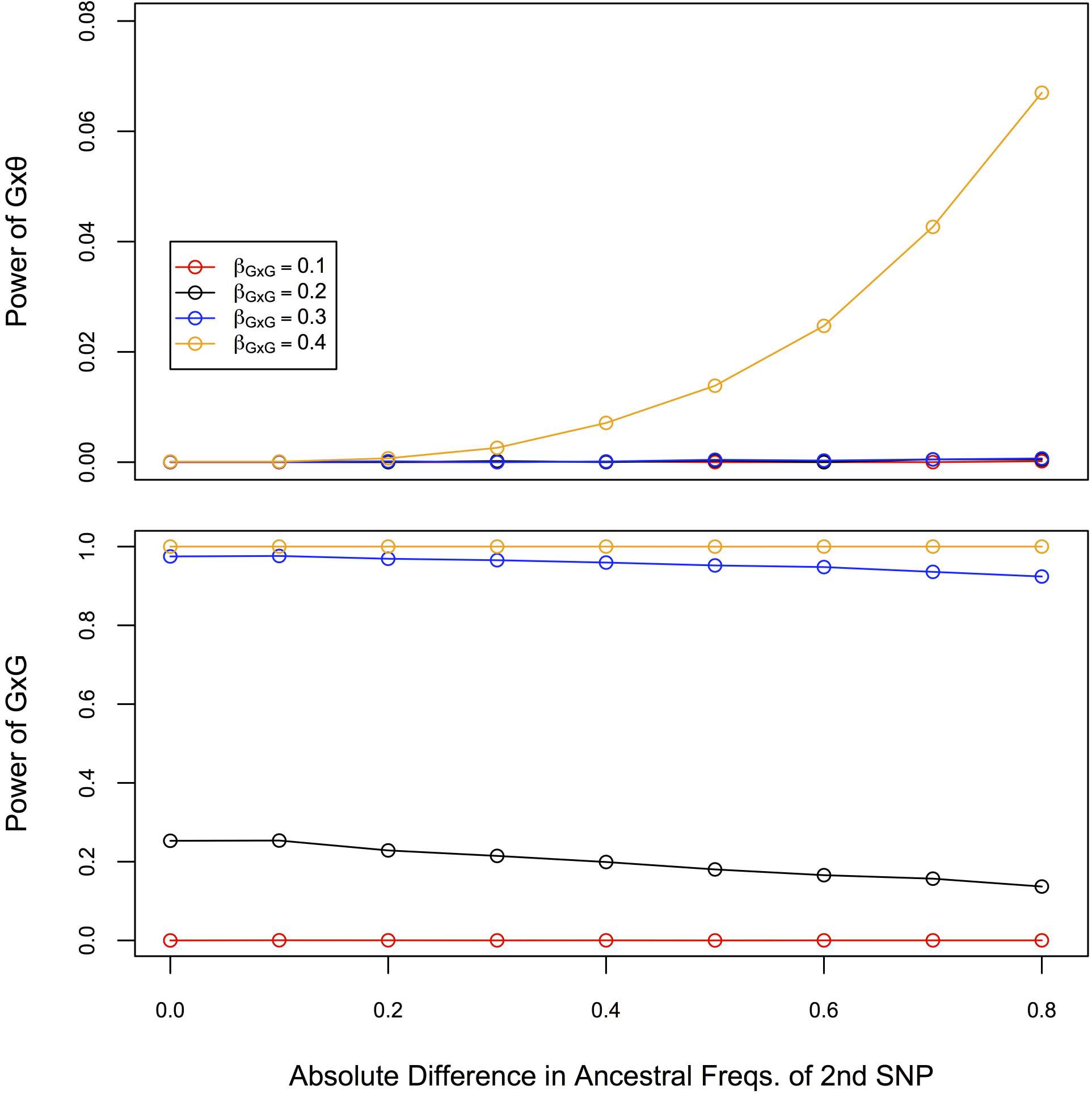
Power Plots for Pairwise Interaction Simulations. Power of testing *G*×*θ* (a) versus testing pairwise SNPs directly (b) as a function of the difference in the ancestral allele frequencies at a differentiated SNP.

However, when multi-way interactions are considered, AITL may become more powerful since differentiated SNPs across the genome will be correlated with genetic ancestry. These simulations are important as other studies have suggested that higher order interactions may be important for some traits[1,30,31]. To evaluate the ability of *θ* to serve as a proxy for multiple (independent) differentiated SNPs, we simulated a scenario where a candidate SNP *z* had interactions with *m* SNPs (see Simulation Framework). For each interaction, we assumed a small interaction effect size (*β*_*G*×*G*_ = 0.025), which would not be detectable using a pairwise approach, as we demonstrated in the pairwise simulation. Figure 2 shows that AITL is better powered to detect the existence of interactions than a pairwise approach in the presence of multiple interacting SNPs with a candidate SNP.

**Figure 2.**
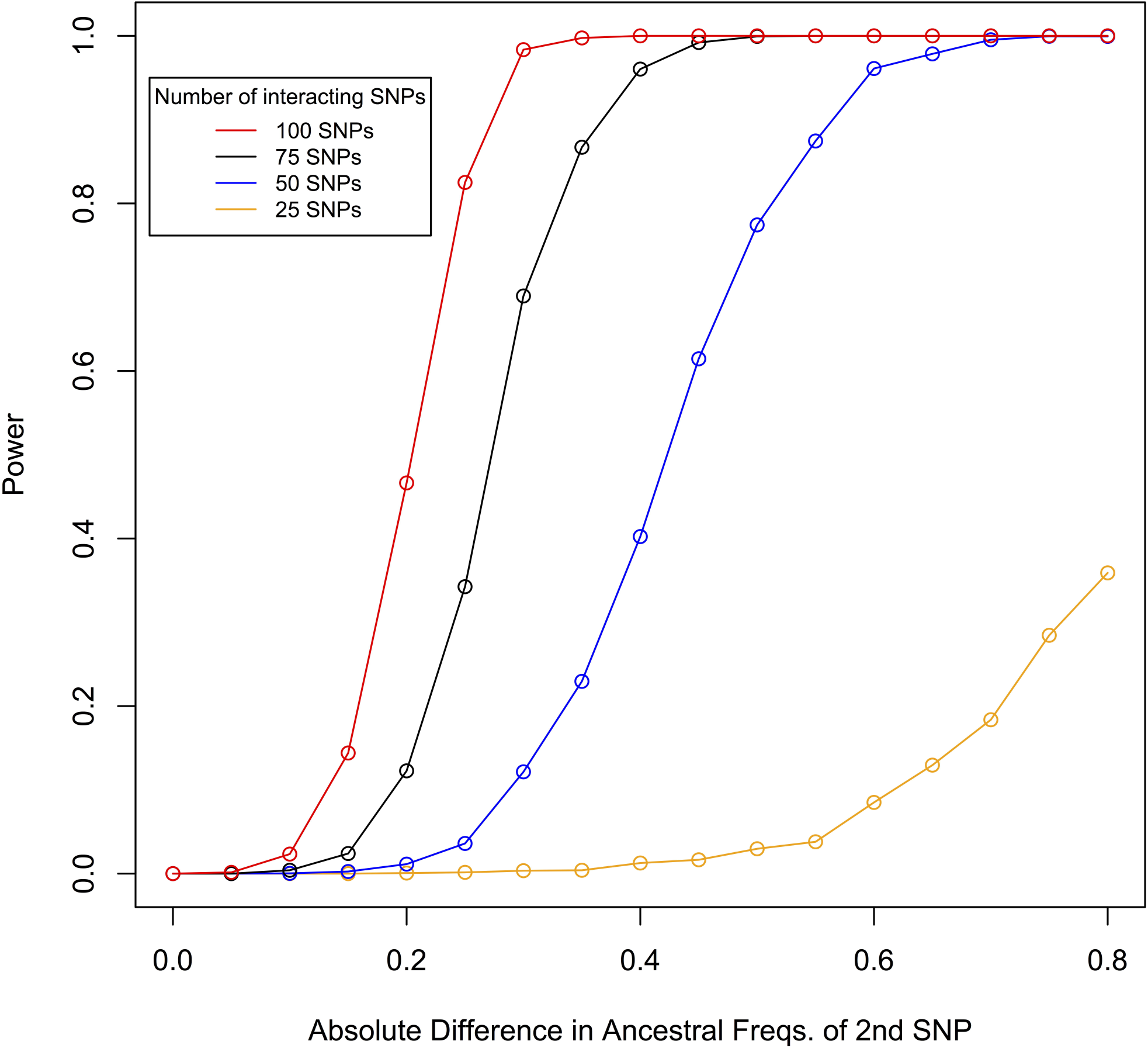
Power Plots for Multi-way Pairwise Interaction Simulations. Power of testing *G*×*θ* as a function of the difference in the ancestral allele frequencies for multiple interacting SNPs.

### Power When Using *θ* as a Proxy Environmental Covariate

When assessing the utility of *θ* as a proxy for an environmental covariate *E*, we simulated 3000 individuals. *E* was simulated such that it was correlated with the global ancestries in varying degrees (see Simulation Framework). Figure 3 shows the power of the AITL as a function of the Pearson correlation between 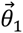and *E*. The power of testing *E* directly is exactly the power of the AITL when the correlation is equal to 1. As expected, as the correlation increases, the power increases as well. When the effect size is 0.1, the power to detect a *G*×*E* interaction is low whether one uses *θ*_1_ or *E*. However, both tests are much better powered for effect sizes greater or equal to 0.2, with the AITL’s power being dependent on the level of correlation. Note that using *θ* as a proxy for *E* is equivalent to testing GxE in the presence of measurement error. Under the assumption of non-differential error with regard to the outcome (e.g. the correlation between *θ* and *E* is equal among cases and control) such a test is underpowered but has a controlled type I error rate under the null[32].

**Figure 3.**
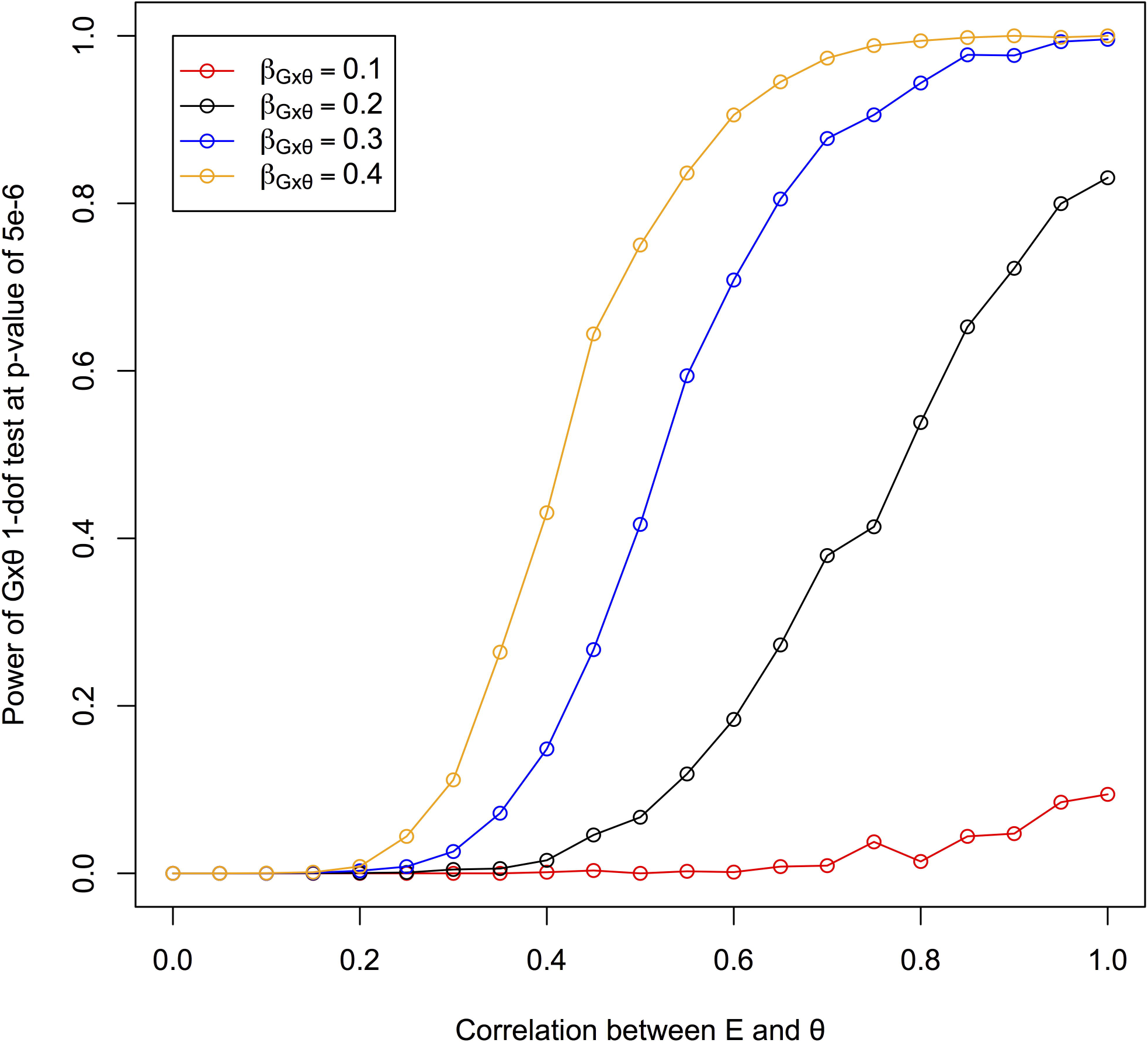
Power Plots for *G*×*E* Interaction Simulations. Power of testing *G*×*θ* as a function of the correlation between an environmental covariate and genetic ancestry.

### Differential LD

To demonstrate that differential LD has the potential to cause inflated test-statistics, we ran 10,000 simulations of 1000 admixed individuals. For each individual we simulated 2 SNPs, a causal SNP and a tag SNP. The LD between the tag SNP and causal SNP was different based on the ancestral background the SNPs were on (see Simulation Framework). Over 10,000 simulations, we computed the mean 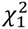 test-statistic for the AIT and the AITL. We note that the phenotypes for these simulations were generated under a model that assumed no interaction. We observed a mean 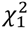 = 0.996 with a standard deviation of 1.53 for AITL. AIT, which does not condition on local ancestry, had a mean 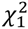 = 3.59 with a standard deviation of 3.60. We also looked at genomic control *λ*_*GC*_, the ratio of the observed median *χ*^2^ over the expected median under the null[33]. *λ*_*GC*_ compares the median observed *χ*^2^ test-statistic versus the true median under the null. In our simulations, we observed *λ*_*GC*_ = 5.81 for AIT and *λ*_*GC*_ = 0.980 for AITL (see Supplementary Figure S1). Last, we computed the proportion of test-statistics that passed a p-value threshold of .05 and .01 in our simulations. The AIT had 3687 statistics passing a p-value of .05 and 1687 at a threshold of .01, whereas AITL had 464 and 96 at the same p-value thresholds. The results for AITL are as expected under a true null. The results from our simulations show that not accounting for local ancestry can result in inflated test-statistics and can potentially lead to false positive findings.

## Real Data

### Coriell Gene Expression Results

We first applied our method to the Coriell gene expression dataset[34]. The Coriell cohort is composed of 94 African-American individuals and the gene expression values of ~8800 genes in lymphoblastoid cell lines (LCLs). Since African Americans derive their genomes from African and European ancestral backgrounds, we tested for interaction between a given SNP and the proportion of European ancestry, *θ*_*EUR*_. Each SNP by *θ*_*EUR*_ term was tested once for association with the expression of the gene closest to the SNP. We observed well-calibrated statistics with a *λ*_*GC*_ equal to 1.04 (see Supplementary Figure S2). In the LCLs, we found that interaction of rs7585465 with *θ*_EUR_ was associated with ERBB4 expression (AITL *p* = 2.95×10^−8^, marginal *p* = 0.404) at a genome-wide significant threshold (*p* ≤ 5×10^−8^). rs7585465 has a ‘C’ allele frequency of 0.218 in the Corriell data and appears to be differentiated between CEU and YRI with allele frequencies of 0.619 and 0.097 in the respective populations.

Given that the gene expression values come from LCLs (all cultured according to the same standards), the SNPs may be interacting with epigenetic alterations due to environmental exposures that have persisted since transformation into LCLs. This scenario is unlikely, and we believe that signals are driven by multi-way epistatic interactions. In our simulations, we showed that using *θ* as a proxy for a single highly differentiated SNP is underpowered compared to testing all pairs of potentially interacting SNPs directly. However, there are many SNPs that are highly differentiated across the genome with which *θ* will be correlated. It is therefore possible that *θ* is capturing the interaction between the aggregate of many differentiated trans-SNPs (i.e. global genetic background) and the candidate SNP. This is consistent with a recently reported finding, conducted in human iPS cell lines, that genetic background accounts for much of the transcriptional variation[2,35].

Although we believe the ERBB4 result to be representative of multi-way epistasis, we performed a standard pairwise interaction test (see Methods) to check for interaction between rs7585465 and other SNPs genome-wide. Interestingly, we found that the standard interaction test (see Methods) showed substantial departure from the null with a *λ*_*GC*_ equal to 1.8 (see Supplementary Figure S3). Since the interaction of rs7585465 by *θ* was significant, the pairwise interaction test-statistics of rs7585465 by any SNP *j* can be inflated if *j* is correlated with *θ*. We found that including the original significant SNP by *θ* term in the null (see Methods) brought the *λ*_*GC*_ down to 1.05, and controlled for such scenarios in this dataset (See Supplementary Figure S3). As we had previously anticipated, identifying the exact interactions driving the SNP by *θ* interaction proved to be difficult. We found one borderline significant SNP (rs4839709, *p* = 3.08×10^−7^) but no interactions that passed genome-wide significance. These results are consistent with what we have observed in simulations, in which even though a standard pairwise interaction test is underpowered to detect interactions, AITL is able to identify the main locus involved in a multi-way interaction.

### GALA II Case-Control

To determine if our method is biased in large structured GWAS data, we applied AITL to case-control data from a study of asthmatic Latino individuals called the Genes-environments and Admixture in Latino Americans (GALA II) [36]. The dataset includes 1158 Mexicans and 1605 Puerto Ricans, which were analyzed separately. Case status was assigned to individuals if they were between the ages of 8 and 40 years with a physician-diagnosed mild to moderate-to-severe asthma. Additionally, they had to have experienced 2 or more asthma related symptoms in the previous 2 years at the time of recruitment[37]. In the Mexicans and Puerto Ricans there were 548 and 797 cases, respectively. In our analysis, we also included BMI, age, and sex as additional covariates. We observed well-calibrated statistics with a *λ*_*GC*_ equal to 1.00 and 0.98 in the Mexicans and Puerto Ricans, respectively (see Supplementary Figure S5). In contrast to the molecular phenotype data, searches for interactions in these phenotypes did not yield any findings passing genome-wide significance. This is consistent with previous disease studies that have failed to find many replicable interactions in disease studies[28]. In the data here, the lack of any findings may be due to the relatively small sample size or because the effects of the interactions are extremely small (if they exist for covariates correlated with *θ*_*EUR*_).

### GALA II Methylation Results

We searched for interactions in methylation data derived from a study of GALA II asthmatic Latino individuals[36]. The methylation data is composed of 141 Mexicans and 184 Puerto Ricans. As the phenotype, we used DNA methylation measurements on ~300,000 markers from peripheral blood. As we had done with gene expression, we tested for interaction between a given SNP and *θ*_*EUR*_ using AITL. All SNPs within a 1 MB window centered around the methylation probe were tested. We used the European component of ancestry because it is the component shared most between Mexicans and Puerto Ricans (see Table 1). We observed well-calibrated test-statistics with *λ*_*GC*_ equal to 1.06 in the Mexicans and 0.96 in the Puerto Ricans (see Supplementary Figure S6). We tested 128,794,325 methylation-SNP pairs, which result in a Bonferroni corrected p-value cutoff of 3.88×10^−10^. However, this cutoff is extremely conservative given the tests are not independent. We therefore report all results that are significant at 5×10^−8^ in either set as an initial filter. We found 5 interactions in the Mexicans and 3 in the Puerto Ricans that are significant at this threshold (see Table 2).

**Table 1.**
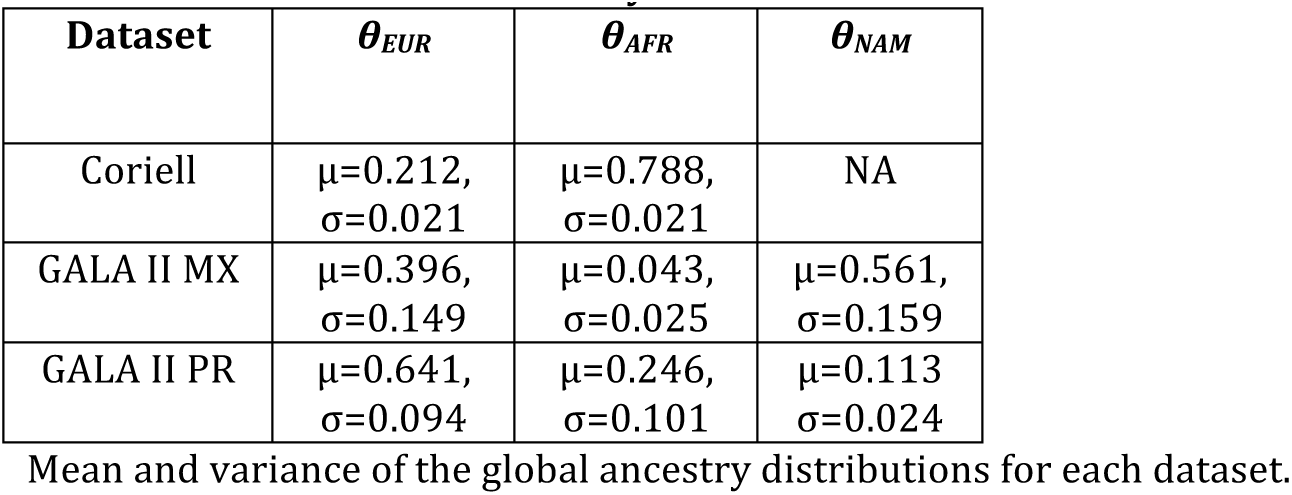
Distribution of Ancestry in Coriell and GALA II.

**Table 2.** GALA II DNA Methylation Analysis Results.

| GALA II Population | Probe Gene | Probe ID | rsid | Distance of SNP to Probe | Marginal p-value | AITL p-value | AITL Replication p-value |
| --- | --- | --- | --- | --- | --- | --- | --- |
| MX | CNFN | cg14327995 | rs16975986 | 280795 | 2.49E-09 | 5.69E-09 | 9.27E-03 |
| MX | C11orf95 | cg16678159 | rs7106153 | 249768 | 2.58E-01 | 2.52E-08 | 9.39E-02 |
| MX | NA | cg05697734 | rs1560919 | 13711 | 1.14E-01 | 2.21E-08 | 8.18E-03 |
| MX | TNK2 | cg01792640 | rs67217828 | 278866 | 4.49E-01 | 6.38E-09 | 1.43E-02 |
| MX | HDAC4 | cg06533788 | rs925736 | 9548 | 4.51E-01 | 3.09E-09 | 2.80E-02 |
| PR | NA | cg07436864* | rs8117083 | 31813 | 7.46E-02 | 1.34E-09 | 5.34E-03 |
| PR | NA | cg16803083* | rs4312379 | 63847 | 3.69E-01 | 2.29E-08 | 2.31E-04 |
| PR | SERPINA6 | cg10025865 | rs17091085 | 247796 | 6.83E-01 | 2.97E-08 | 8.05E-03 |

P-values for AITL applied to the methylation data in the GALA II Latinos. MX and PR denote Mexicans and Puerto Ricans respectively in the GALA II population columns. The probe gene column shows the gene that the methylation probe lies in. The marginal column is the p-value for standard linear regression of methylation on genotype while controlling for population structure. \* indicates results that replicated between the Mexicans and Puerto Ricans.

Unlike the Coriell individuals, who are 2-way admixed, the GALA II Latinos are 3-way admixed and derive their ancestries from European, African, and Native American ancestral groups. Consequently, to confirm that incomplete modeling or better tagging on one of the non-European ancestries was not driving the results, we retested all significant interactions including a second component of ancestry for AITL. In the case of the Mexicans, we included African and European ancestry, and in the case of the Puerto Ricans, we included European and Native American ancestry. Even after adjusting for the second ancestry the interactions between SNP and *θ*_*EUR*_ remained highly significant (see Supplementary Table 1).

As we did for the gene expression data, we attempted to identify pairwise interactions involved in the methylation data results. For each genome-wide significant result, we performed a standard pairwise interaction test of all SNPs with the original SNP found to be significant with AITL. We were unable to identify any significant interactions after applying genomic control to the results. For all tests, we included the significant SNP by *θ* term (see Methods) in the null. For this dataset, unlike the gene expression data, we observed substantial remaining departure from the null (see Supplementary Table S2) even after including the original significant SNP by *θ* term, suggesting there may be other factors that need to be accounted for when testing for interactions in admixed populations. The results from our pairwise scan are what we would anticipate, given that in simulations only AITL (not the standard pairwise interaction test) was able to identify the main locus involved in the multi-way interaction.

We then performed a replication study of the significant Puerto Rican associations in the Mexican cohort and vice versa. To account for the fact that we are replicating eight total results across both populations, we used a Bonferroni corrected p-value threshold equal to .05/8 = 6.25×10^−3^. The interaction of rs4312379 and rs4312379 with ancestry in the Puerto Ricans replicated in the Mexicans. Furthermore, there was a highly significant enrichment of low p-values in the replication study among the discovery results (permutation *p* < 1×10^−4^). Furthermore, 5 out of the 6 non-replicating results have a p-value less than 0.05 (binomial test *p* < 1.8×10^−6^). The results of the permutation and binomial test suggests that the interactions that did not replicate are likely to do so with bigger sample sizes. It is important to note that replicated interactions and the enrichment for low p-values do not necessarily indicate that the same genetic or environmental covariates are interacting with the genetic locus in both populations. The covariates correlated with *θ*_EUR_ in one population are not necessarily those correlated with *θ*_*EUR*_ in the other population. There may be correlations which exist in both populations but *θ*_*EUR*_ serves as a proxy for all such correlated covariates and therefore should not be necessarily viewed as a proxy for any specific one. Overall, our results from the GALA II (methylation) cohort suggest there are both genetic and environmental variables contributing to epistasis that have yet to be discovered in admixed individuals.

## IV Discussion and Conclusions

For many disease architectures, interactions are believed to be a major component of missing heritability[38]. Finding new interactions has proven to be difficult for logistical, statistical, biological, and computational reasons. In this study, we have demonstrated that in admixed populations, testing for *G*×*θ* interactions can be leveraged to overcome some of the difficulties typically encountered when searching for interactions. The computational cost is minimal and has the same order as running a standard GWAS.

One drawback of our method is that it does not identify which covariate is interacting with a genetic locus. Nevertheless, the approach can show whether an interaction effect exists in a given dataset and if it does exist, our method ensures that an underlying genetic or environmental covariate(s) is correlated with ancestry. Additionally, in the case where there is no marginal effect, our approach identifies new loci and shows that the genetic locus influences the phenotype and exerts its effects through interactions, which has important implications for the genetic architecture of the phenotype. The relative contribution of additive and non-additive genetic effects to variability in molecular phenotypes and disease risk is an important area of investigation, and our approach provides a direct test for detecting non-additive contributions[39].

Environmental covariates are often not consistently measured across cohorts whereas genetic ancestry is nearly perfectly replicable. Testing for the presence of interaction using a nearly perfectly reproducible covariate may enhance our understanding of the genetic basis of disease and other traits. Our method also provides the additional benefit of not being confounded by interactions between unaccounted-for covariates[40].

Association testing for interaction effects involving continuous environmental exposures in the context of mixed-models remains an open problem. For binary environmental exposures, it has been shown that mixed-models control for population structure nominally better than including genetic ancestry (or principal components) as a covariate[41]. Because it is unclear how mixed-models perform with continuous environmental exposures, especially those correlated with ancestry, in our analyses we took the standard approach of filtering related individuals and including ancestry as a covariate.

It has been shown that 2-step analyses may be more powerful for detecting interactions when exposures are binary [42–44]. However, these studies have primarily been done in a single homogeneous population, and the correct null distribution for the interaction effect must assume that the 2^nd^ stage procedure is independent of the marginal effect test-statistic. In real data, using a 2-step approach in conjunction with AITL to test for interactions may be problematic because the interaction effect size will not necessarily be independent of the marginal effect size, as the allele frequency at any SNP will be a function of ancestry in an admixed population. Additionally, only 1 of the interaction results that we report here had a marginal effect (p< 0.05) and thus would have been missed by a 2-step approach. Thus, our approach can serve to complement or extend the frequently used 2-step procedure for detecting interaction effects.

Results from our multi-way epistasis simulation analyses and empirical data in cell lines suggest that genetic ancestry is a good proxy for genetic background, since all highly differentiated SNPs across the genome will be correlated with genetic ancestry. Our simulations also demonstrated that genetic ancestry can be a good proxy for an environmental covariate depending on the correlation between the two. However, it may be the case that there are multiple environmental factors interacting with a genetic locus, all of which are correlated with *θ* in differing degrees and effect sizes. Such a situation would mirror what we saw in our multi-way *G*×*G* simulations where a single interaction may not be detectable by using a traditional *G*×*E* test, but because *θ* aggregates the effects of all interacting covariates, AITL would be able to detect it. There are also other contexts in which modeling SNP by *θ* may be useful, such as using variance components. For example, SNP by *θ* interaction terms can be used in a mixed-model framework to test for interaction effects because genetic ancestry is correlated with many genetic markers and environmental covariates[45].

For some traits, there may be systematic differences between ancestral populations in the genetic effects on the trait. In admixed individuals with these ancestral populations, the effect of genetic variation on phenotype will be reflected in the correlation between phenotype and *θ*, thereby affecting epistatic and *G*×*E* interactions. It will be interesting to see how much of the phenotype-ancestry correlations are due to epistatic and *G*×*E* interactions.

In our analysis of real data, we discovered gene by *θ* interactions associated with genes that have known interactions. In the GALA II Mexicans, the interaction of rs925736 with ancestry was associated with the methylation of HDAC4, a known histone deaceytlase (HDAC). In concert with DNA methylases, HDACs function to regulate gene expression by altering chromatin state[46]. In Europeans, HDACs have been shown to be associated with lung function through direct genetic effects and through environmental interactions[47,48]. For the GALA II Puerto Ricans, rs17091085 showed an interaction associated with the methylation state of SERPINA6. Of note, interaction between birth weight and SERPINA6 has been previously associated with Hypothalamic-Pituitary-Adrenal axis function[49]. Further investigations of our interaction findings are thus warranted.

In the GALA II (methylation) dataset, two of the eight significant associations replicated and, in general, the results had an enrichment of low p-values in the replication dataset. However, we note that if the interactions detected by AITL are multi-way epistasis it is more likely that the results will replicate. This is because most SNPs differentiated in the Mexicans will still be differentiated in the Puerto Ricans, and thus still be correlated with *θ*. If the interactions detected by AITL are *G*×*E* interactions, then the interactions are less likely to replicate because the same environmental covariate(s) will need to be correlated with ancestry in both groups.

Another caveat is that the Mexicans and Puerto Ricans, though independent, are part of the same study and occasionally technical artifacts, such as issues with genotyping or measuring methylation, can affect downstream analyses of both populations. For our analyses, we have taken careful quality-control steps to ensure that this is not the case and there is no apparent inflation of test-statistics as demonstrated by our values for genomic control. Future research of interactions using AITL should keep such caveats in mind.

We investigated in detail the potential of single SNP-SNP interactions driving the results that were found both in the gene expression and methylation datasets. As demonstrated by the wide range of *λ*_*GC*_ values, we observed that non-linear effects can cause substantial departure from the null when testing for pairwise SNP-SNP interactions. This is especially true when testing for interaction between SNPs *s* and *j*, where *s* has a significant interaction with *θ* and *j* is correlated covariates that are also correlated with *θ*. As we saw in the gene expression data, including the significant SNP by *θ* term can properly control for such situations, but its use in standard pairwise interaction tests warrants further investigation.

Our analysis revealed the existence of interactions but does not provide a direct way to determine the covariate that is interacting with a SNP. Further methodological work is required to uncover the exact environmental exposures or genetic loci with which SNPs are interacting. The existence of gene by *θ* interactions in GALA II underscores why modeling interactions should be considered for future association studies and for heritability estimation in admixed populations.

## V. Materials and Methods

Our approach is best illustrated with an example. First consider testing a SNP s for interaction with an environmental covariate *E*. *θ* can serve as a proxy for *E* if the two are correlated, even if *E* is unknown or unmeasured (see Figure 4a). Now consider testing s for interaction with a SNP *j*≠*s* that is highly differentiated in terms of ancestral allele frequencies. For example, a SNP that has a high allele frequency in one ancestral population and a low allele frequency in the other ancestral population. *θ* can be used as a proxy for *j* because *θ* and the genotypes of SNP *j* will be correlated. Consider the case where *j* has a frequency of 0.9 in population 1 and frequency of 0.1 in population 2. Individuals with large values of *θ*_1_ are more likely to have derived *j* from population 1 and on average have greater genotype values at *j*. Similarly, individuals with small values of *θ*_1_ are more likely to have derived *j* from population 2 and on average have smaller genotype values. Thus, *θ* will be correlated with the genotypes of the individuals for highly differentiated SNPs and can serve as a proxy for detecting interactions (see Figure 4b).

**Figure 4.**
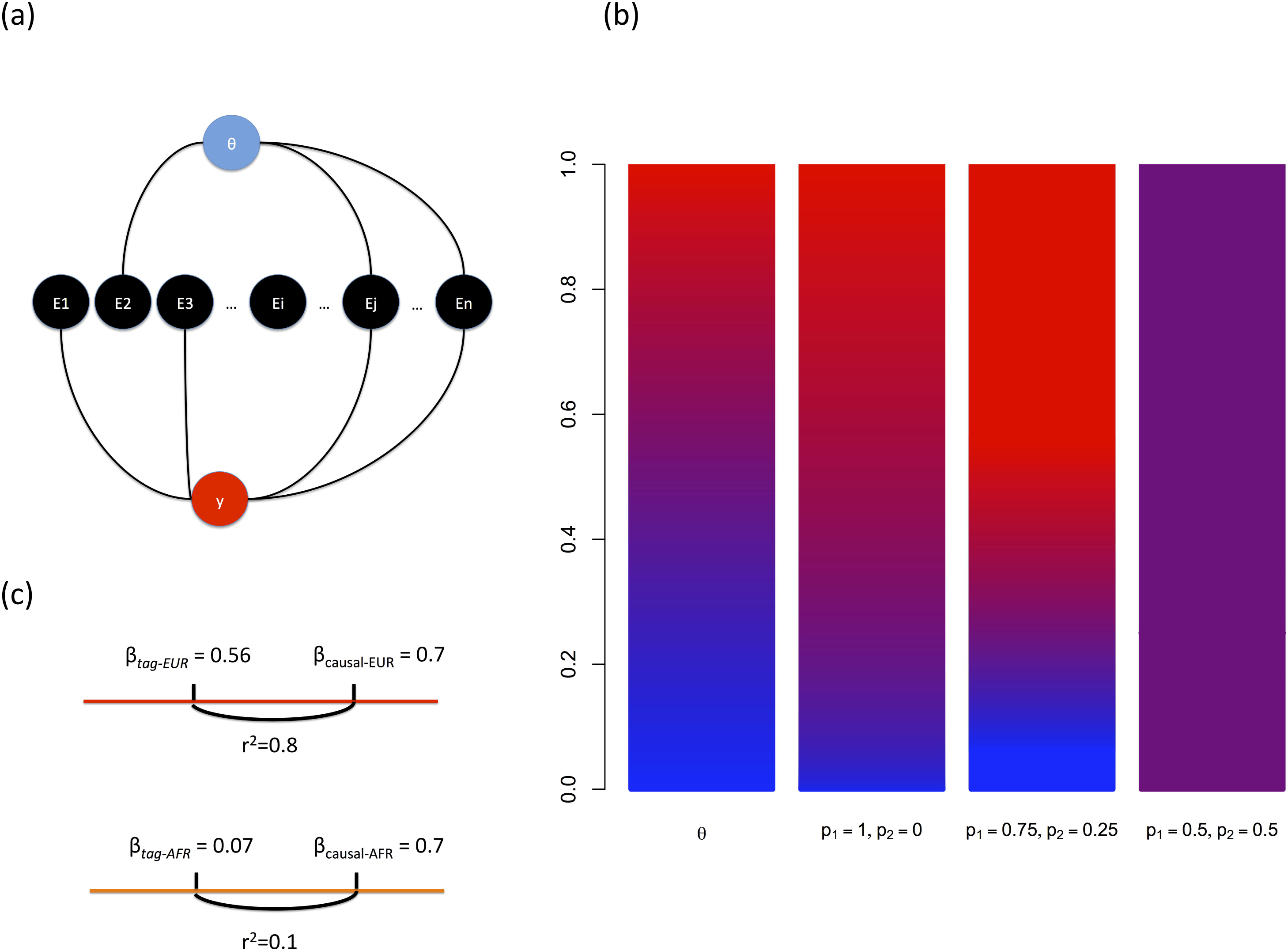
Examples of How Genetic Ancestry Can Be A Proxy for Interacting Covariates. (a) Model of how genetic ancestry *θ* can be correlated with various environmental exposures, some of which affect a phenotype. (b) Example of how the correlation between the probability of an AA genotype (bars 2-4) and values of *θ* (bar 1) increase with higher levels of SNP allele frequency differentiation. In this plot p_1_ and p_2_ denote the allele frequency of allele A in ancestral populations 1 and 2 respectively. (c) Example of how effect sizes at a tag-SNP may differ due to differential LD on distinct ancestral backgrounds (here, EUR and AFR).

Consider an admixed individual *i* who derives his or her genome from *k* ancestral populations. We denote individual *i*’s global ancestry proportion as *θ*_*i*_ = 〈*θ*_*i*1_, *θ*_*i*2_, ,*θ*_*ik*_〉, where Σ_*k*_ *θ*_*ik*_ = 1. The local ancestry of individual *i* at a SNP *s* is denoted as *γ*_*ais*_ ∈ {0,1, 2} and is equal to the number of alleles from ancestry *a* ∈ {1 … *k*} inherited at SNP *s*. Current methods allow us to estimate ancestry directly from genotype data both globally and at specific SNPs[9,50,51]. We denote the genotype of an individual *i* at SNP *s* as *g*_*is*_ ∈ {0,1,2} and the corresponding phenotype as *y*_*i*_.

In this work, we model continuous phenotypes in an additive linear regression framework. Assuming *n* (unrelated) individuals, define 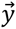 to be the vector of all individuals’ phenotypes. The model for the phenotype is then

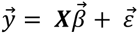

where 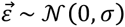 is a *n*×1 vector of error terms, ***X*** is a *n*×*v* matrix of *v* covariates, and 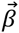 is a *v*×*1* vector of the covariate effect sizes. We note that in our notation 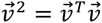 for a vector 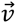. Assuming independence, the likelihood under this model is:

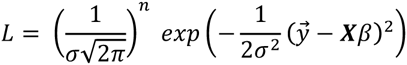

We can compute the log-likelihood ratio statistic (D) using a maximum likelihood approach:

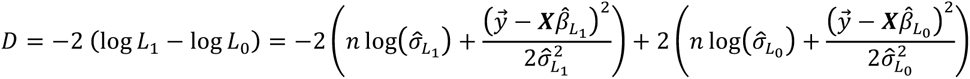

We note that for a case-control phenotype we would use the following likelihood and log-likelihood ratio statistic:

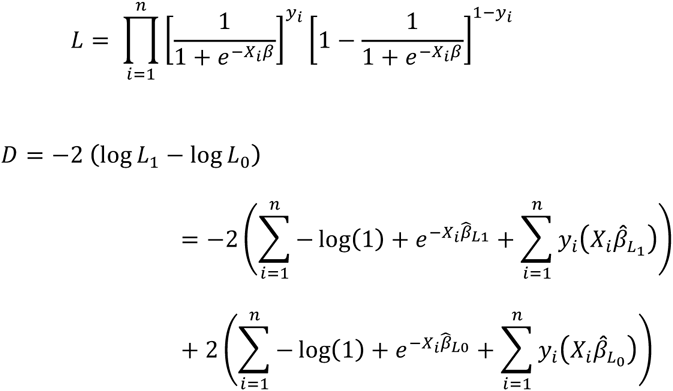

where *X*_*i*_ is the *i*-th row of the matrix ***X***, which correspond to the covariates of individual *i*.

For linear regression, the maximum likelihood estimator (MLE) of the effect sizes is 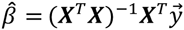, and the MLE of the error variance is 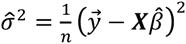. Here, *L*_1_ is the likelihood under the alternative and *L*_0_ is the likelihood under the null. 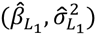 and 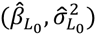 are the effect sizes and error variance estimates that maximize the respective likelihoods. *D* is distributed as *χ*^2^ with *k* degrees of freedom (*df*), where *k* is the number of parameters constrained under the null.

### 1-df Ancestry Interaction Test (AIT)

The first test we present is the standard direct test of interaction. We test for a SNP’s interaction with *θ* instead of an environmental covariate or another genotype. Let 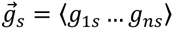 be the vector of the individuals’ genotypes at SNP *s*, 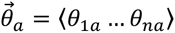) be the vector of their global ancestries for ancestry *a*, and 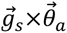 be the vector of interaction terms which result from the component-wise multiplication of the genotype and global ancestry vectors. We test the alternative hypothesis (*β*̂_*G*×*θ*_ ≠ 0) against the null hypothesis (*β*̂_*G*×*θ*_ ≠ 0).

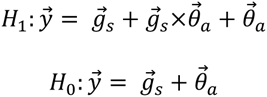

In this test of interaction, we test a single ancestry versus the other ancestries that may be present in the population of interest. One parameter is constrained under the null which results in a statistic with *k*=1 *df*. Let *β*̂_*L*_{0, 1}_(*S*)_, *β*̂_*L*_{0, 1}_(*G*×*θ*)_, and *β*̂_*L*_{0, 1}_(*θ*)_ denote the effect sizes of genotype, interaction, and global ancestry under a given hypothesis respectively. The statistic is given below.

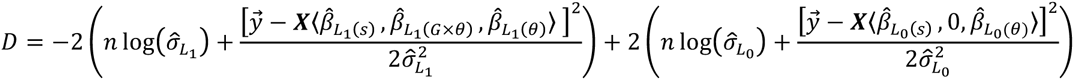

where *X* is an *n*×3 matrix composed of 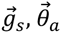, and 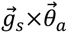 as columns.

### 1-df Ancestry Interaction Test with Local Ancestry (AITL)

Given that the individuals we analyze in this work are assumed to be admixed, there is potential for confounding due to differential LD. An interaction that is not driven by biology could occur due to the possibility that a causal variant may be better tagged by a SNP being tested on one ancestral background versus another (See Figure 4c). We account for the different LD patterns on varying ancestral backgrounds by including local ancestry as an additional covariate in AITL. By including local ancestry, we assume that the SNP being tested is on the same local ancestry block as the causal SNP that it may be tagging. Such an assumption is reasonable because admixture in populations such as Latinos and African Americans are relatively recent events and their genomes have not undergone many recombination events. As a result, local ancestry blocks on average stretch for several hundred kilobases[52,53].

Let 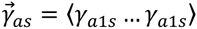 be the vector of local ancestry calls for all individuals for ancestry a and let 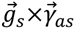 be the interaction terms from piecewise multiplication of the two vectors. We use the following alternative and null hypotheses.

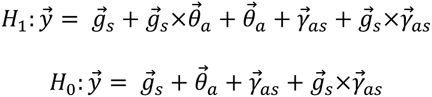

Here we are testing for an interaction effect, i.e. *β*̂_*G*×*θ*_ ≠ 0, and constrain one parameter under the null resulting in a statistic with *k*=1 *df*. Let *β*̂_*L*_{0, 1}_(*G*×*γ*)_ and *β*̂_*L*_{0, 1}_(*γ*)_ denote the effect sizes of the interaction between genotype and local ancestry and just local ancestry, respectively. The log likelihood ratio statistic is given by

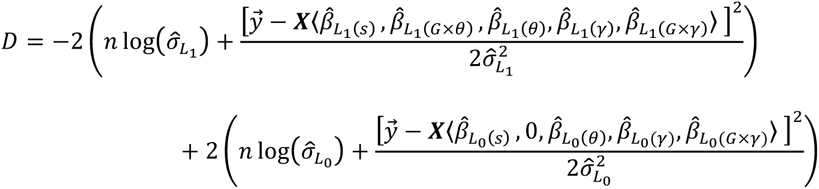

where ***X*** is an *n*×5 matrix composed of 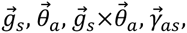, and 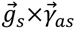 as columns. All of these test-statistics are straightforwardly modified to jointly incorporate several ancestries in the case of multiway admixed populations.

### Standard Pairwise Test of Interaction and Controlling Confounding in Admixed Populations

Here we present the standard approach for testing for interaction between two SNPs. We use the following alternative and null hypotheses.

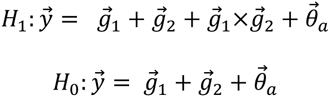

If AITL is significant for a given SNP *s*, then any SNP *j* tested for interaction with *s* may be biased if *j* is correlated with covariates that are also correlated with *θ*. Furthermore, if the effects of the covariates correlated with *θ* are non-linear then controlling for the main effects of the SNPs and ancestry will account for the non-linear effects. We thus, propose the use of the following alternative and null hypotheses.

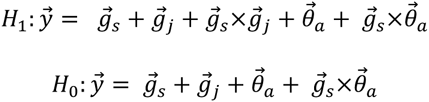

We note that the utility of this test will require further investigation (see Discussion).

### Simulation Framework

For all our simulations, we simulated 2-way admixed individuals. Global ancestry for ancestral population 1 (*θ*_l_) was drawn from a normal distribution with *μ*. = 0.7 and *σ* = 0.2. Individuals with *θ*_l_ 1 or *θ*_1_ < 0 were assigned a value of 1 or 0, respectively. We simulated phenotypes of individuals to investigate our method in four different scenarios: *G*×*E* interactions, pairwise epistatic interactions, multi-way epistatic interactions, and false positive interactions due to local differential tagging.

To simulate phenotypes under the situation of a *G*×*E* interaction, we simulated a single SNP. For each individual *i*, we assigned the local ancestry or the number of alleles derived from population 1 (*γ*_*ai*_) for each haplotype by performing two binomial trials with the probability of success equal to *θ*_*i*1_. We then drew ancestry specific allele frequencies following the Balding-Nichols model by assuming a *F*_*ST*_ = 0.16 and drawing two ancestral frequencies, *p*_1_ and *p*_2_, from the following beta distribution[54].

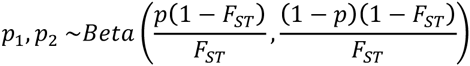

where *p* is the underlying MAF in the entire population and is set to 0.2. Genotypes were drawn using a binomial trial for each local ancestry haplotype with the probability of success equal to *p*_1_ or *p*_2_ for values of *γ*_*ai*_ = 0 or 1, respectively. Environmental covariates correlated with *θ*_1_, *E*_*i*_, were generated for each individual *i* by drawing from a normal distribution ***N***(*μ* = *θi*1, *σ*_*E*_), where *σ*_*E*_ is the standard deviation of the environmental covariates. *σ*_*E*_ was varied from 0 to 5 in increments of 0.005 to create 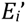 that were correlated with individuals’ global ancestries in varying degrees. We generated phenotypes for individuals assuming only an interaction effect by drawing from a normal distribution, ***N***(*μ* = *β*_*G*×*E*_ × *g*_*i*1_×*E*_*i*_, *σ* = 1) for a given interaction effect size(*β*_*G*×*E*_).

To simulate phenotypes based on pairwise epistatic interactions, we simulated two SNPs. At both SNPs, we assigned the local ancestry values as described for the *G*×*E* case. We assigned genotypes for individuals at the first SNP assuming an allele frequency of 0.5 for both populations and drawing from two binomial trials. We assigned genotypes at the second SNP over a wide range of ancestry specific allele frequencies to simulate different levels of SNP differentiation. Ancestry specific allele frequencies were initially *p*_1_ = *p*_2_ = 0.5 and iteratively increasing *p*_*1*_ by 0.005 while simultaneously decreasing *p*_*2*_ by 0.005 until *p*_*1*_ = 0.05 and *p*_*2*_ = 0.95. Genotypes at the second SNP were drawn using the same approach described for *G*×*E*. Using the simulated genotypes, phenotypes were drawn from a normal distribution, ***N***(*μ* = *β*_*G*×*G*_×*g*_*i*1_×*g*_*i*2_,*σ* = 1), where *g*_*is*_ is the genotype for individual *i* at the simulated SNP *s*.

To simulate phenotypes based on multi-way epistatic interactions, we simulated a SNP *z* and *m* (independent) SNPs with pairwise interactions with *z*. Genotypes for individuals at SNP *z* were assigned assuming an allele frequency of 0.5 for both populations and drawing from two binomial trials. Genotypes at the *m* interacting SNPs were assigned in the same manner as the 2^nd^ SNP in the pairwise interaction simulations. Using the simulated genotypes, phenotypes were drawn from a normal distribution, 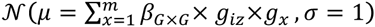 where *g*_*is*_ is the genotype for individual *i* at the simulated SNP *s*.

To simulate the scenario of differential LD on different ancestral backgrounds leading to false positives, we simulated phenotypes based on a single causal SNP that was tagged by another SNP. At both SNPs, local ancestries were assigned as described previously and genotypes were drawn using ancestry specific allele frequencies. Ancestral allele frequencies were assigned such that the average *r*^*2*^ between the causal and tag SNP was 0.272 on the background of ancestral population 1 and 0.024 on the background of ancestral population 2. Thus, the tag SNP was only a tag on the population1 background and not on the population 2 background. Phenotypes were drawn from a normal distribution, ***N***(*μ* = *β*_*Causal*_×*g*_*ic*_, *σ* = 1), assuming no interaction and *β*_*Causal*_ = 0.7, where *g*_*ic*_ is the genotype of individual *i* at the causal variant.

We implemented our approach in an R package (GxTheta), which is available for download at http://www.scandb.org/newinterface/GxTheta.html

## Ancestry Inference

Global ancestry inference was done using ADMIXTURE [9] and local ancestry inference was done using LAMP-LD [55]. CEU and YRI from 1000 Genomes Phase 3 [56] were used as the European and African reference panels. For the Native American reference panels, 95 Native Americans genotyped on the Axiom LAT1 array were used[57].

## Filtering for Related Individuals

All analyses in real data were filtered for related individuals due to the possibility of cryptic relatedness causing false positives. To filter for related individuals, we estimated kinship coefficients between all pairs of individuals using REAP [58]. We defined two individuals as related if they had a kinship coefficient greater than 0.025. For a pair of related individuals, we removed the one with a greater number of other individuals to whom he or she was related. In the case of a tie, we removed one of the pair at random.

## Data Normalization

### Gene Expression Normalization

Gene expression data (see Results) were first standardized for each gene such that mean expression was 0 and variance was 1. We then computed a covariance matrix of individual’s expression values and performed PCA on the covariance matrix. Residuals were computed for all expression values by adjusting for the top 10 principal components and the mean for each gene was added back to the residuals. Due to the high dynamic range of gene expression compared to methylation we conservatively chose to additionally perform quantile normalization. We then sorted the gene expression residuals and used the quantiles of their rank order to draw new expression values from a normal distribution, ***N***(*μ* = 0, *σ* = 1), by using the inverse cumulative density function^24,25^.

### Methylation Data Normalization

Raw methylation values (see Results) were first normalized using Illumina's control probe scaling procedures. All probes with median methylation less than 1% or greater than 99% were removed and the remaining probes were logit-transformed as previously described[59]. To control for extreme outliers, we truncated the distribution of methylation values. For a given probe, we first computed the mean and standard deviation of the methylation values. We then set any methylation values deviating more than 2.58 standard deviations from the mean to the methylation value corresponding to the 99.5^th^ quantile.

## Availability of Supporting Data

The Coriell data is available from dbGAP under accession number phs000211.v1.p1. The GALA and SAGE data is available by emailing the study organizers at https://pharm.ucsf.edu/gala/contact.

## Authors’ Contributions

DSP, IE, EK, EE, EH and NZ designed research. DSP, IE, EK, ERG, and NZ performed research. DSP, IE, EK, EE, CE, CRG, JMG, EG, HA, CJY, EE, EH, and NZ contributed new reagents/analytic tools. DSP, ERG, and NZ wrote the manuscript. All authors read and approved the final manuscript.

## Description of Additional Data Files

The following data are available with the online version of this paper. The Supplemental contains QQ-plots for the simulations and real analyses performed as well as a table containing p-values for the 2-component ancestry analysis of the GALA methylation data.

## Acknowledgements

We would like to thank Lancelote Leong for his helpful manuscript comments.

